# SwrA extends DegU over an UP element to activate flagellar gene expression in *Bacillus subtilis*

**DOI:** 10.1101/2023.08.04.552067

**Authors:** Ayushi Mishra, Anna C. Hughes, Jeremy D. Amon, David Z. Rudner, Xindan Wang, Daniel B. Kearns

## Abstract

SwrA activates flagellar gene expression in *Bacillus subtilis* to increase the frequency of motile cells in liquid and elevate flagellar density to enable swarming over solid surfaces. Here we use ChIP-seq to show that SwrA interacts with many sites on the chromosome in a manner that depends on the response regulator DegU. We identify a DegU-specific inverted repeat DNA sequence and show that SwrA synergizes with phosphorylation to increase DegU DNA binding affinity. We further show that SwrA increases the size of the DegU footprint expanding the region bound by DegU towards the promoter. The location of the DegU inverted repeat was critical and moving the binding site closer to the promoter impaired transcription more that could be explained by deactivation. We conclude that SwrA/DegU forms a heteromeric complex that enables both remote binding and interaction between the activator and RNA polymerase in the context of an interceding UP element. We speculate that multimeric activators that resolve cis-element spatial conflicts are common in bacteria and likely act on flagellar biosynthesis loci and other long operons of other multi-subunit complexes.

**IMPORTANCE:** In Bacteria, the sigma subunit of RNA polymerase recognizes specific DNA sequences called promoters that determine where gene transcription begins. Some promoters also have sequences immediately upstream called an UP element that is bound by the alpha subunit of RNA polymerase and is often necessary for transcription. Finally, promoters may be activated by transcription factors that bind DNA specific sequences and help recruit RNA polymerase to weak promoter elements. Here we show that the promoter for the 32 gene long flagellar operon in *Bacillus subtilis* requires an UP element and is activated by a heteromeric transcription factor of DegU and SwrA. Our evidence suggests that SwrA oligomerizes DegU over the DNA to allow RNA polymerase to interact with DegU and the UP element simultaneously. Heteromeric activator complexes are known but poorly-understood in bacteria and we speculate they may be needed to resolve spatial conflicts in the DNA sequence.

## INTRODUCTION

Bacterial flagella are complex, multi-subunit, trans-envelope machines that propel cells to swim in liquid or swarm over solid surfaces. Flagella are composed of dozens of different subunits, expressed in a series of hierarchical stages roughly corresponding to the order in which the subunits are assembled (1–4). The top of the hierarchy is often a transcription factor called a master regulator that directly enhances the expression of the earliest genes in flagellar synthesis. Mutation of the master regulator tends to impair or abolish motility while over-expression of the master regulator can lead to hyper-flagellation and, in some systems, promote swarming motility (5). Master regulators bind upstream of one or more promoters in the flagellar regulon and often require the formation of heteromeric, multiprotein complexes (6–8). While clearly important for motility gene expression, these master regulators are among the most species-specific and the least-studied components of the hierarchy.

In *Bacillus subtilis,* the master flagellar regulator is SwrA, a small (117 amino acid), positively-charged protein that is narrowly-conserved in a closely-related subset of species within the genus *Bacillus* (9,10). First discovered as a factor required for swarming but not swimming motility (9), SwrA activates the constitutive *P_flache_* promoter of the 32-gene long *fla/che* operon encoding early-stage flagellar biosynthesis components, over a relatively narrow (∼4-fold) range (11–14). Moderate SwrA levels support a high frequency of swimming cells in liquid and SwrA levels increase on a surface for enhanced flagellar synthesis to support swarming motility (13; 15-17). The mechanism by which SwrA enhances *P_flache_* promoter activity is poorly-understood but SwrA has been shown interact with the DNA binding protein DegU (14, 18-20).

DegU is a response regulator that functions as a transcription factor. Cells lacking DegU have pleiotropic phenotypes that include defects in genetic competence, exoprotease production, exopolymer production, biofilm formation, and motility (21–30). A consensus binding sequence for DegU has not been determined and the mechanism by which DegU differentially regulates the wide variety of targets under its control is poorly-understood (31,32). DegU binds DNA in both phosphorylated and unphosporylated forms and each may bind targets differently (14,25,33-35). DegU is phosphorylated either by its cognate soluble kinase DegS (36,37) or by the small metabolite acetyl phosphate (38) and two small proteins DegQ and DegR enhance the phosphorylated state (25,39). Finally, SwrA binds to the DegU receiver domain and may alter DegU’s DNA binding activity (19,20). Consistent with being a modulator of DegU function, cells lacking SwrA have phenotypes beyond defects in motility, many of which overlap with cells lacking DegU (40–42).

Here we characterized SwrA association with the chromosome using Chromatin immunoprecipitation coupled to deep sequencing (ChIP-seq) and found that SwrA is enriched at DNA sequences in a manner that depends on DegU. SwrA enhanced DegU binding at a variety of target sites *in vitro* with the highest affinity displayed at *P_flache_*, consistent with genetic results suggesting that it was the primary biological SwrA target (13). Despite extensive sequence analysis, no SwrA or DegU consensus site was detected bioinformatically, but a perfect 5-8-5 inverted repeat within *P_flache_* was shown to be required for SwrA/DegU binding and activity. SwrA interaction synergized with phosphorylation to increase DegU’s DNA binding affinity and expanded the DegU binding site likely by inducing DegU oligomerization. Inverted repeat location was critical and moving the repeat closer to the promoter abolished promoter activity by disrupting a cis-acting UP element. Thus we suggest that DegU oligomerization induced by SwrA is necessary to allow both the remote binding of DegU and interaction with RNA polymerase at the promoter. Complex heteromeric activators may be generally required for promoters with UP elements, and we speculate that a number of poorly-understood transcriptional modulators may act enabling the transcription factor to contact RNA polymerase without interfering with, or otherwise occluding, UP elements.

## RESULTS

### SwrA interacts with DNA indirectly

SwrA is the master regulator of motility in *B. subtilis,* and SwrA activates the *P_flache_* promoter that controls 32 genes involved in flagellar assembly and chemotaxis (13,14). Other targets of SwrA have been reported but the extent of the SwrA regulon, and the mechanism of SwrA-mediated transcriptional activation are poorly-understood (13,19,20,41,42). To investigate whether SwrA associates with DNA *in vivo*, we performed chromatin immunoprecipitation coupled to deep sequencing (ChIP-Seq) on wild type and *swrA* mutant cells. Cultures were treated with formaldehyde, and after lysis and DNA fragmentation, SwrA was immunoprecipitated with antibodies raised against the full-length protein. After reversing the crosslinks, the DNA associated with SwrA was subjected to next generation sequencing. Chromatin immuno-enrichment was calculated as the ratio of ChIP-Seq signal to genomic DNA plotted as peaks in 1 kb windows that spanned the entire genome. SwrA candidate binding sites were defined as peaks that were enriched in wild type replicates but not in the *swrA* mutant control (**Fig 1A**).

**Figure legend 1:**
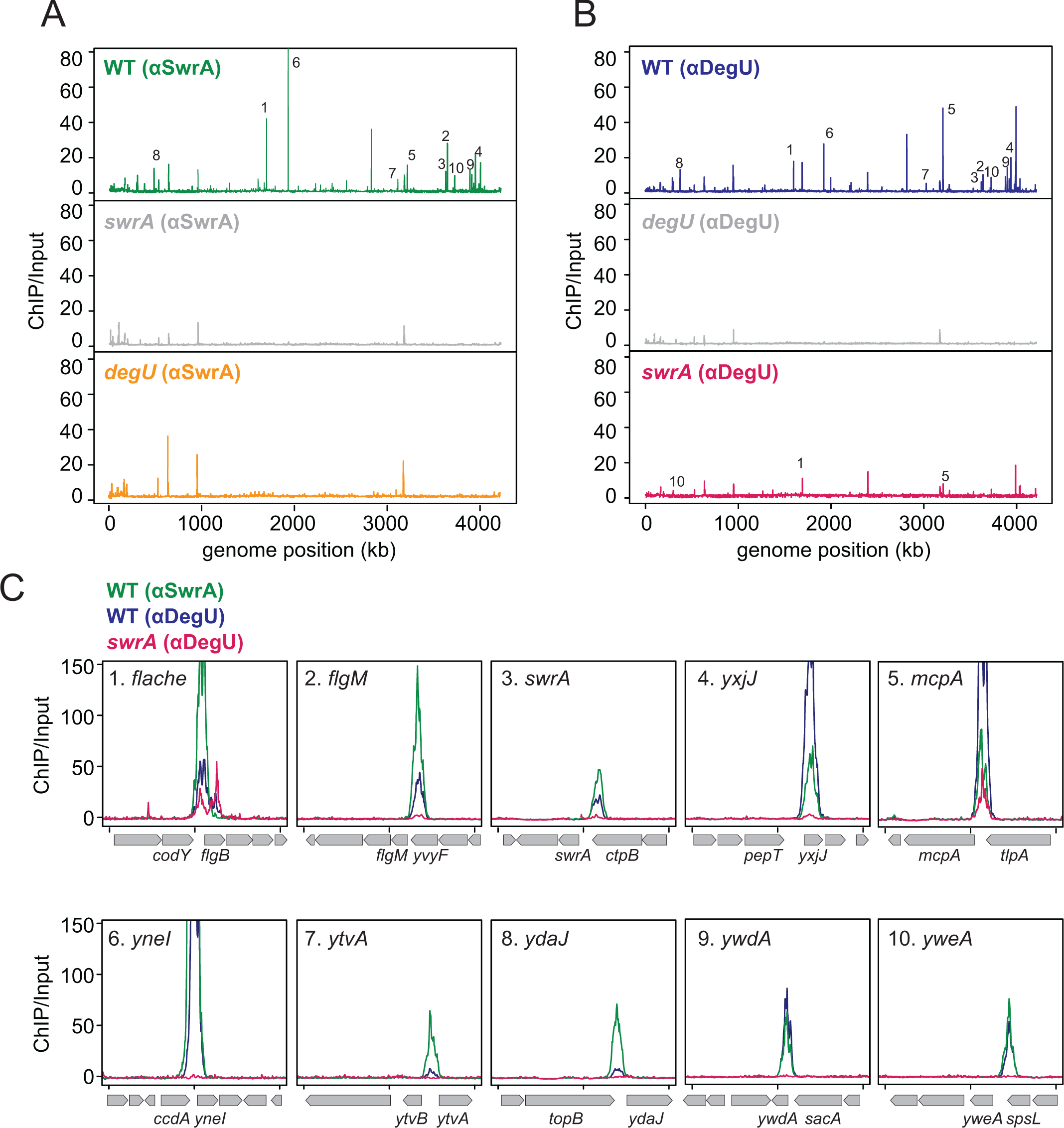
SwrA enriches a subset of DegU-enriched promoters. A) ChIP-Seq analysis using a primary antibody to SwrA (αSwrA). The number of sequencing reads were normalized by the total number of reads for each sample. The fold of enrichment (ChIP/Input) were calculated and plotted in 1 kb bins. The following strains were used to generate this panel: WT (3610), *ΔswrA* (DS2415), and *ΔdegU* (DS3649). Peaks of particular interest were numbered 1-10. B) ChIP-Seq analysis using a primary antibody to DegU (αDegU). Data were processed in the same way as in panel A. The following strains were used to generate this panel: WT (3610), *ΔdegU* (DS3649) and *ΔswrA* (DS2415). Peaks of particular interest were numbered 1-10 to match the same peaks in panel A. C) ChIP-Seq analysis of peaks 1-10 from panels A and B enlarged to show detail over a 4 kb range. Each panel is named according to the promoter region for the likely gene downstream. Gene size and identity is indicated below the X-axis. The X-axis is marked in 2 kb intervals. Green lines indicated WT ChIP-Seq data using αSwrA antibody, blue lines indicate WT ChIP-Seq data using αDegU antibody, and magenta lines indicate *swrA* mutant ChIP-seq data using αDegU antibody.

SwrA was enrichment at thirty-four genomic locations and all but two were located in intergenic regions consistent with possible promoters (**Table S1**). To investigate whether SwrA affected transcription of the neighboring genes, 13 promoter regions containing strongly-enriched peaks (*P_flache_*, P*_flgM_*, P*_swrA_*, P*_yxjJ_*, P*_mcpA_*, *P_ycdC_*, P*_yneI_, P_ytvA_, P_ydaJ_, P_ywdA_, P_yweA_, P_tlpA_*, and *P_sacX_)* were cloned upstream of the *lacZ* gene encoding β-galactosidase, and β-galactosidase activity was measured in various genetic backgrounds. As anticipated, expression of *P_flache_-lacZ* decreased relative to wild type in cells lacking SwrA, and increased when was overexpressed, consistent with previous reports (**Fig 2A**) (13,14). The remaining reporters responded to the presence and absence of SwrA in a variety of ways. Expression from four promoters, *P_flgM_*, *P_swrA_*, *P_yxjJ_*, and *P_mcpA_* was reduced ∼2-fold in a *swrA* deletion and increased ∼2-fold when *swrA* was artificially overexpressed in a manner that was similar to *P_flache_* (**Fig 2A**). Five promoters, P*_yneI_, P_ytvA_, P_ydaJ_, P_ywdA_,* and *P_yweA_* produced low but detectable levels of LacZ activity that was not altered by either mutation or overexpression of *swrA* (**Fig 2A**). Finally, three promoters, *P_tlpA,_ P_ycdC_* and *P_sacX_*, produced no activity above background (<2 Miller units, MU) in any strain and were omitted from the study. We conclude that while ChIP-Seq indicated SwrA-enrichment of several promoter regions, enrichment did not necessarily reflect an effect of SwrA on reporter expression. We further conclude that SwrA either directly or indirectly activated the P*_fla/che_*, P*_flgM_*, *P_swrA_*, P*_yxjJ_*, and P*_mcpA_* promoters.

**Figure legend 2:**
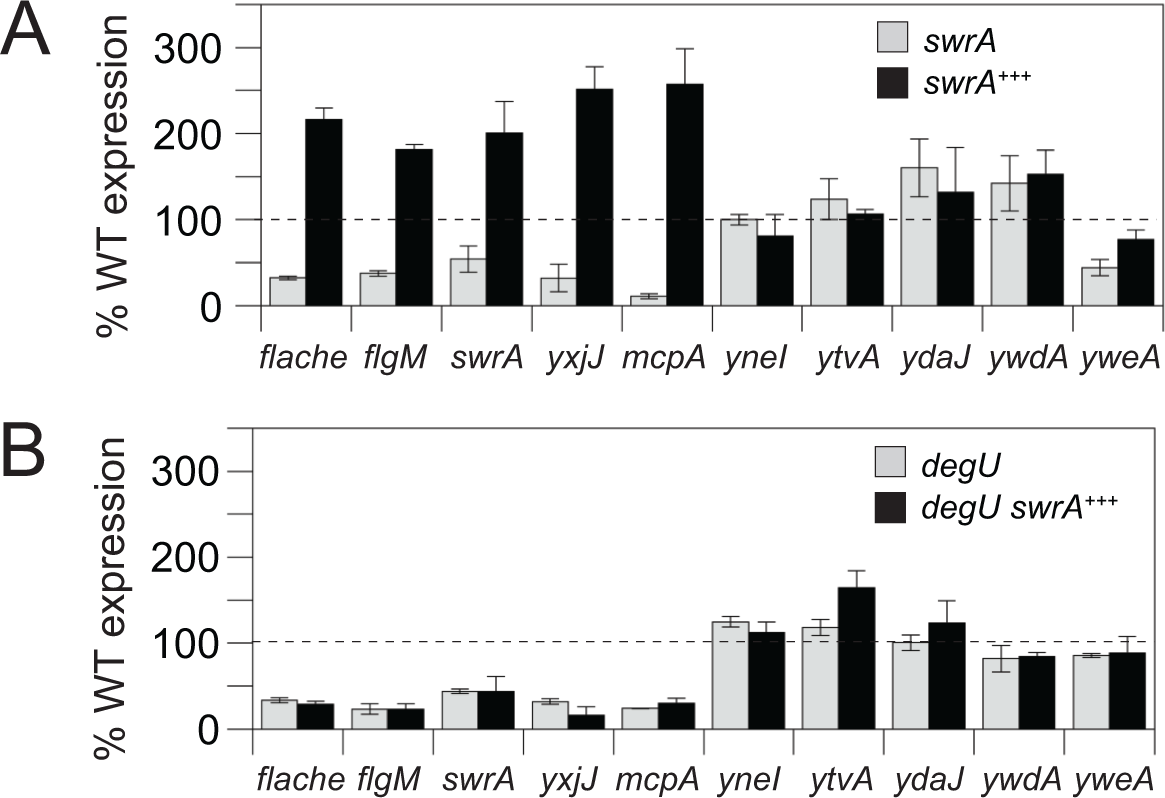
SwrA/DegU activate expression from a subset of enriched promoters. A) β-galactosidase activity from the indicated promoter region fused to the *lacZ* gene. Light gray bars indicate expression in a *swrA* mutant and black bars indicate expression when *swrA* was overexpressed from an IPTG-inducible promoter in meroploid. Activity was normalized to 100% wild type expression (dashed line). Error bars are the standard deviation of three replicates. The following strains were used to generate this panel: *P_flache_-lacZ* (DK4730, DK4918), *P_flgM_-lacZ* (DK4870, DK4919), *P_swrA_-lacZ* (DK6624, DK6625), *P_yxjJ_-lacZ* (DK4733, DK4929), *P_mcpA_-lacZ* (DK4732, DK4928), *P_yneI_-lacZ* (DK4731, DK4927), *P_ytvA_-lacZ* (DK6500, DK6051), *P_ydaJ_-lacZ* (DK6473, DK6477) *P_ywdA_-lacZ* (DK6474, DK6478), *P_yweA_-lacZ* (DK6476, DK6480). B) β-galactosidase activity from the indicated promoter region fused to the *lacZ* gene. Light gray bars indicate expression in a *degU* mutant and black bars indicate expression when swrA was overexpressed from an IPTG-inducible promoter in meroploid. Activity was normalized to 100% wild type expression (dashed line). Error bars are the standard deviation of three replicates. The following strains were used to generate this panel: *P_flache_-lacZ* (DK4734, DK4979), *P_flgM_-lacZ* (DS3103, DK4983), *P_swrA_-lacZ* (DK6626, DK6650), *P_yxjJ_-lacZ* (DK4737, DK4982), *P_mcpA_-lacZ* (DK4736, DK4981), *P_yneI_-lacZ* (DK4735, DK4908), *P_ytvA_-lacZ* (DK6502, DK6550), *P_ydaJ_-lacZ* (DK6481, DK6509) *P_ywdA_-lacZ* (DK6482, DK6524) *P_yweA_-lacZ* (DK6484, DK6510). Raw data included in table S5.

One way in which SwrA could enrich target promoter regions is by binding directly to DNA. To determine whether SwrA bound DNA directly, electrophoretic mobility shift assays (EMSA) were performed on seven different promoter regions. The *P_flache_* promoter was chosen as a known SwrA target and *P_flgM_*, *P_yxjJ_*, *P_swrA_*, and *P_yneI_* were added as candidates from the present study. Two negative controls were included: the promoter *P_comK_* expressing the gene for the master activator of competence gene expression ComK, and the promoter *P_hag_* expressing the gene for the flagellar filament protein Hag. The seven different promoter fragments were PCR amplified, radiolabeled, incubated with purified SwrA protein at a range of protein concentrations, resolved by native gel electrophoresis and analyzed by phosphorimager. In each case, addition of SwrA failed to alter migration of the radiolabeled DNA fragment except at the highest concentration of SwrA added (1 μM) (**Fig S1**). These data argue that SwrA is not sufficient for DNA binding.

### SwrA-DNA interaction is DegU-dependent

Current evidence suggests that SwrA interacts with the *P_flache_* promoter indirectly by interacting with the response regulator DegU (19,20). Consistent a requirement of both proteins for DNA binding, cells lacking either SwrA or DegU are defective in swarming motility even when the remaining protein was artificially overexpressed (**Fig S2**) (9,20,25,26,43). To test whether DegU is required for SwrA’s association with DNA, we repeated the SwrA ChIP-Seq in a *degU* mutant strain. In the absence of DegU, all SwrA-dependent peaks were abolished (**Fig 1A**). Furthermore, the *degU* mutation reduced the expression of the SwrA target promoters *P_flache_*, *P_flgM_*, *P_swrA_*, *P_yxjJ_*, and *P_mcpA_* to levels similar to that observed in a *swrA* mutant, and SwrA overexpression failed to activate these promoters (**Fig 2B**). We conclude that SwrA association with DNA and transcriptional activation of SwrA-responsive promoters are entirely dependent on the presence of DegU.

To investigate whether SwrA alters DegU DNA binding specificity, we performed ChIP-seq using a DegU antibody. DegU was enriched at 46 locations, many of which were upstream of genes involved in motility, competence, biofilm formation as well as genes of unknown function (**Fig 1B; Table S1**) (44). All 34 SwrA-enriched regions were also enriched in the DegU ChIP-seq (**Fig 1C, Fig S3A**). Furthermore, in cells lacking SwrA, there was a general reduction enrichment in the DegU ChIP-seq (**Fig 1B,1C**). Peaks were sorted into three different DegU-ChIP classes depending on the effect of SwrA on enrichment (**Table S1**). Class I targets (61%) were SwrA-dependent as they were abolished in the absence of SwrA. Class II targets (11%) were SwrA-enhanced as they were reduced but not abolished when SwrA was absent. Class III targets (28%) were SwrA-independent as they appeared unaffected by its absence. We conclude that three-quarters of the DegU-enriched promoters were either enhanced by, or fully dependent on, the presence of SwrA. Therefore, SwrA potentiates binding of DegU to a subset of promoters in its regulon.

In an effort to determine a consensus binding site for DegU, 200 base pair fragments surrounding each ChIP-seq peak center were compiled and subjected to MEME sequence pattern analysis (45). Combining sequences from all DegU peaks indicated an enriched sequence that did not contain a repeat element as might be expected for a response regulator DNA binding sequence (**Fig S4**). Separate analysis of the class I, class II and class III DegU ChIP peaks, provided similarly-enriched sequences (**Fig S4**). We wondered whether our data set might be incomplete due to the absence of peaks located near the characterized DegU-regulated promoters: *P_aprE_* and *P_pgsB_* directing alkaline protease and poly-γ-glutamate synthesis respectively (**Fig S3B**) (27,41,46-48). Accordingly, we performed a DegU ChIP-seq by immune-precipitating DegU from a strain overexpressing the small phosphor-enhancer protein DegQ and a strain expressing the hyper-active DegU allele, DegU^hy32^ (21,25,33). Both the DegQ overproduction (**Fig S5A, Table S2**) and DegU^hy32^ (**Fig S5B, Table S3**) strains identified additional peaks that were SwrA-enhanced, but MEME analysis of each dataset still produced an asymmetric DegU target sequence similar to that of the wild type (**Fig S4**). Moreover, sequence analysis did not indicate how SwrA was differentiating a subset of the DegU peaks for enrichment and there appeared to be no SwrA-specifying sequence in the DNA.

### SwrA increases DegU affinity for DNA and expands the DegU binding site

To better understand the mechanism of SwrA-mediated DegU activation, DegU EMSAs were conducted on the same series of promoters previously used to test for direct interaction by SwrA. DegU bound poorly to each promoter but an electrophoretic mobility shift was observed with *P_flache_* when DegU was phosphorylated by ATP and its cognate kinase, DegS (**Fig S1**) (14,25). Thus, the *P_flache_* promoter seemed to contain the highest affinity binding site of those tested, and we note that previously studied promoters like *P_flgM_* and *P_comK_*, bound by DegU-P and DegU respectively, require higher concentrations than used here (34,49). Next, the concentration of either DegU or DegU-P was held constant and increasing amounts of SwrA were added to the reaction. The presence of SwrA caused a supershift of the *P_flache_* promoter and did so at lower concentrations when DegU was phosphorylated (**Fig 3**). Moreover, the presence of SwrA induced a shift of all promoters except the non-predicted target *P_hag_*, suggesting that SwrA generally enhanced the affinity of DegU-P for its targets.

**Figure legend 3:**
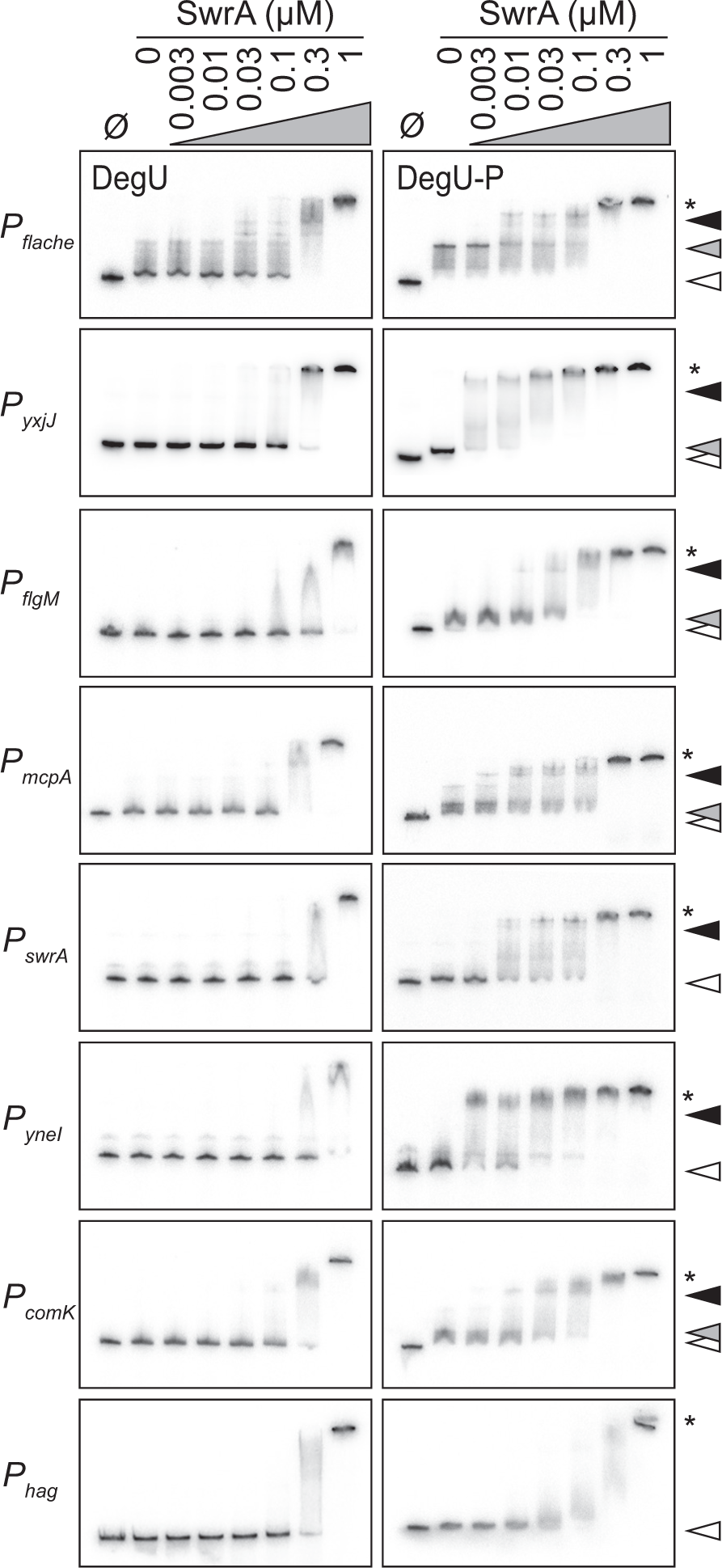
SwrA causes a supershift in DegU DNA binding. Electrophoretic mobility shift assays (EMSA) with the indicated radiolabeled promoter fragment and protein. Ø indicates that no protein was added. Left panels, an increasing about of SwrA was added to a constant 0.3 μM of DegU. Right panels, an increasing amount of SwrA was added to a constant 0.3μM of DegU-P phosphorylated by DegS and ATP. Gray carets indicate the position of DNA shifted by the presence of either DegU or DegU-P alone. Black carets indicated the position of DNA supershifed by the presence of either DegU or DegU-P and SwrA. * indicates aggregation.

To explore the effect of SwrA on DegU DNA binding activity, the concentration of SwrA was held constant, and EMSA experiments were conducted in triplicate on *P_flache_* with increasing amounts of DegU and DegU-P. Addition of SwrA caused diffuse supershifted bands making it difficult to quantify the densitometry of the bound state. Therefore, we calculated the fraction of the unbound state and subtracted the value from 1. Phosphorylation of DegU increased DNA binding affinity by 10-fold (**Fig 4A,B**). SwrA increased the binding affinity of DegU (**Fig 4A**) and DegU-P (**Fig 4B**) by approximately 10-fold and 4-fold respectively, suggesting that SwrA affinity enhancement was phosphorylation independent (**Fig 4**). High levels of SwrA caused smearing of the band signal even for non-target promoters like *P_hag_* but did so with a K_d_ much higher than observed for *P_flache_* (**Fig 4C**). While ChIP-seq is non-quantitative, we note SwrA-enhancement of DegU DNA binding is consistent with the *in vivo* global reduction in DegU enrichment in the absence of SwrA (**Fig 1B**). We conclude that SwrA interacts with DegU and synergizes with phosphorylation to enhance DNA binding.

**Figure legend 4:**
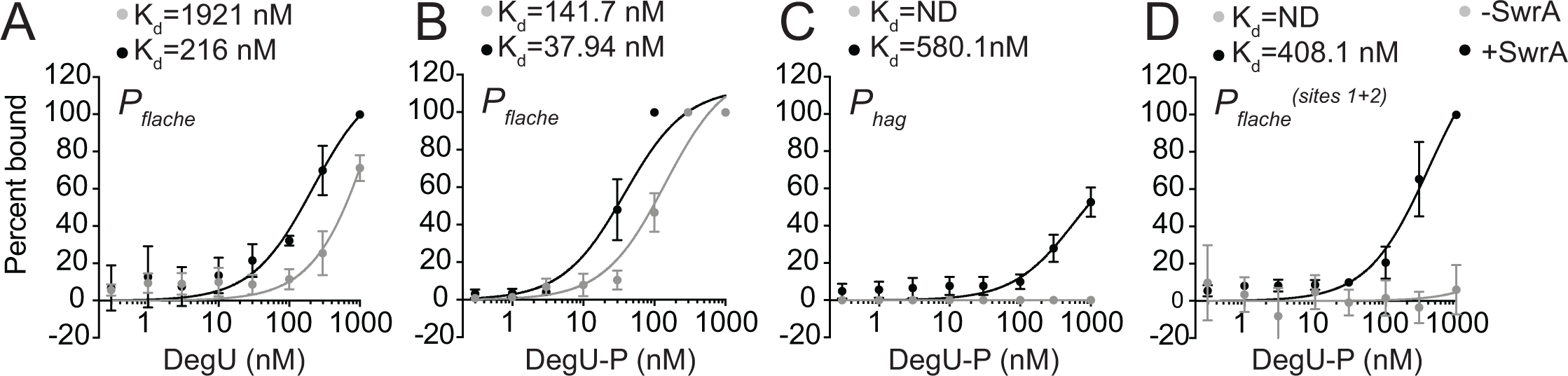
SwrA increases DegU DNA-binding affinity. Densitometry analysis of electrophoretic mobility shift assays (EMSA) (**Fig S9**) with the indicated radiolabeled promoter fragment and various concentrations of either DegU or DegU-P phosphorylated by ATP and DegS. Gray circles are EMSA densitometry in the presence of the indicated form of DegU. Black circles are EMSA densitometry in the presence of the indicated form of DegU and 0.2 μM SwrA. Each point is the average of three replicates and error bars are standard deviations. A) Densitometry values of radiolabeled *P_flache_* promoter sequence shifted by DegU; B) densitometry values of radiolabeled *P_flache_* promoter sequence shifted by DegU-P; C) densitometry values of radiolabeled *P_hag_* promoter sequence shifted by DegU-P; D) densitometry values of radiolabeled *P_flache_* promoter sequence doubly mutated at site 1 and site 2 shifted by DegU-P.

To further explore the effect of SwrA on DegU-P, DNAse I protection assays were conducted on the *P_flache_* (**Fig 5A**) and *P_yxjJ_* (**Fig 5B**) promoters. DegU-P protected a region of the *P_flache_* promoter 40 base pairs (bp) upstream of the −35 box (**Fig 5A**). Centered within the protected region was a perfect 5-8-5 inverted repeat of CTAGG separated by an intervening 8 base pairs (**Fig 5C**). Addition of SwrA to the reaction expanded the protection to both the left and right of the repeat thereby increasing the total protected area to approximately 70 bp. DegU-P alone did not provide DNAse I protection of *P_yxjJ_*at the concentration used, and while the location of the *P_yxjJ_*promoter is unknown, addition of SwrA induced protection again with a width of approximately 70 bp (**Fig 5B**). We conclude that SwrA alters the way in which DegU-P binds DNA and creates a wide footprint of DNase I protection at multiple promoters.

**Figure legend 5:**
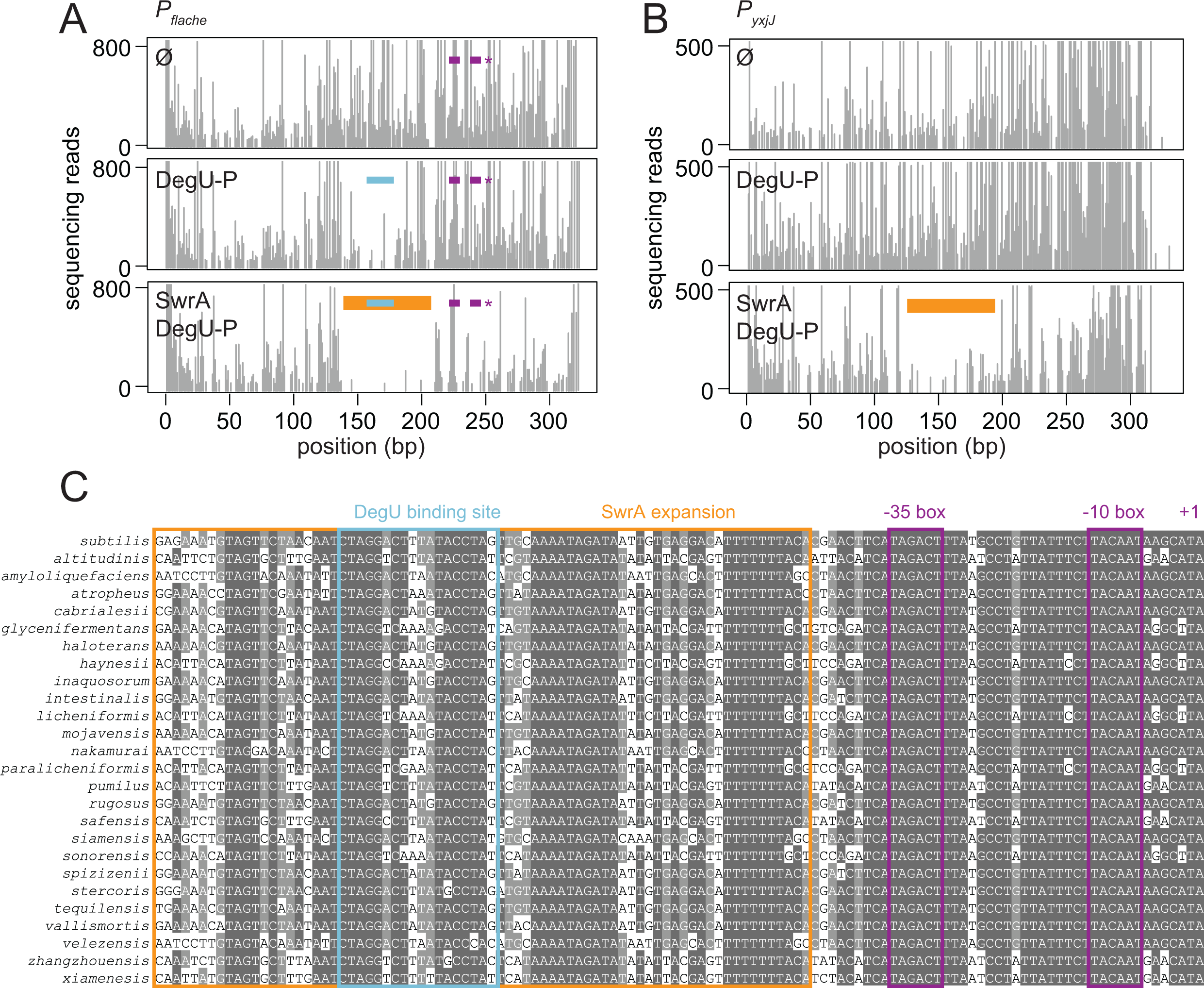
SwrA increases the length of DNA protected by DegU-P. A) DNaseI sequencing footprint analysis of the *P_flache_* promoter. 300 bp fragments of dsDNA were fluorescently labeled on the forward strand, the indicated protein was added, followed by partial digestion with DNase I and sequencing of the digested fragments. Top panel, 1 uM BSA added; middle panel 3 μM of DegU-P added; bottom panel 3 μM of DegU-P and 1μ M of SwrA added. Blue bars indicate the location of the 5-8-5 inverted repeat; orange bars indicate the expanded region of SwrA/DegU-P protection; purple bars indicate the −35 element, the −10 element, and +1 transcriptional start sites of the *P_flache_*promoter are marked with an asterisk. Peaks indicate the number of sequence reads terminating at that location on one strand. B) DNaseI sequencing footprint analysis of the *P_yxjJ_* promoter. Top panel, 1 uM BSA added; middle panel 3 μM of DegU-P added; bottom panel 3 μM of DegU-P and 1 μM of SwrA added. Peaks indicate the number of sequence reads terminating at that location. C) Alignment of the *P_flache_* promoter fragment. Blue boxes highlights the location of the 5-8-5 inverted repeat (repeat sequences in bold); Orange box highlight the expanded region of SwrA/DegU-P protection; purple highlights that −35 and −10 elements of the *P_flache_* promoter and the +1 transcriptional start site.

The consensus sequence to which DegU-P binds is poorly-understood and we focused our attention on the putative 5-8-5 inverted repeat within the *P_flache_*promoter. To test whether the inverted repeat element was important for SwrA/DegU-dependent activation, two bases were changed in each half site of the repeat separately and together at the native site in the chromosome. Mutation of either the promoter distal site (site 1) or the promoter proximal site (site 2) had little effect on swarming motility but mutation of both sites simultaneously caused a severe defect (**Fig 6A**). Overexpression of SwrA shortened the lag period of the single mutants and restored partial swarming to the double mutant. Moreover, EMSA indicated that mutation of both sites simultaneously abolished and dramatically reduced binding of DegU-P in the absence and presence of SwrA respectively (**Fig 4D**). Finally, consistent with an important regulatory element, the sequence upstream of the *P_flache_* promoter, including the putative 5-8-5 repeat, was highly-conserved in a wide variety of *B. subtilis* relatives that also encoded SwrA and DegU (**Fig 5C**) and conservation degraded farther upstream (**Fig S6**). We conclude that the conserved inverted repeat protected in the DegU-P DNase I protection assay is required for SwrA/DegU-dependent activation of *P_flache_*.

**Figure legend 6:**
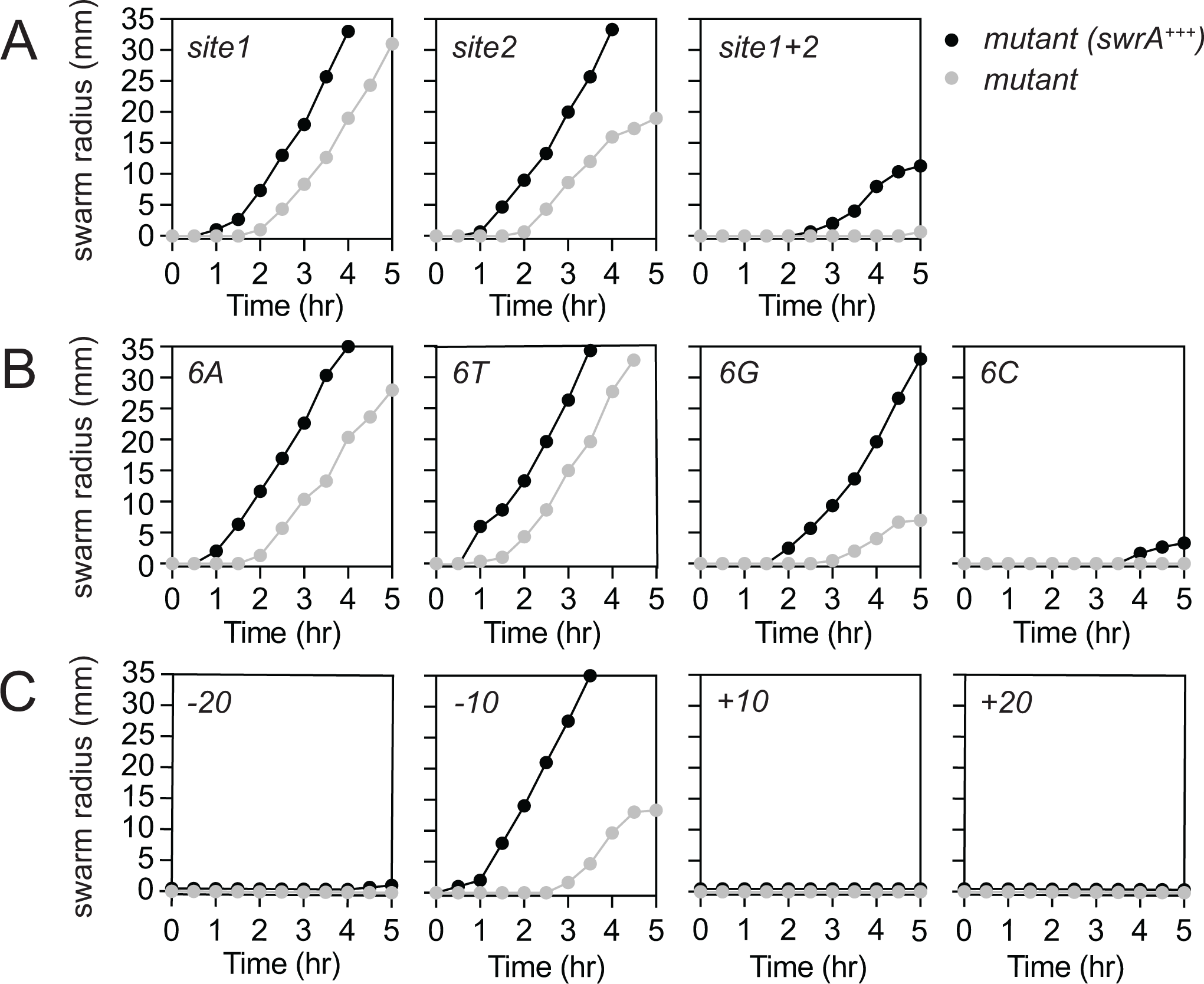
The distance between the DegU binding site and the *P_flache_* promoter is important for activation. A) Quantitative swarm expansion assays of strains mutated for the DegU binding site inverted repeat sequences (gray circles), and strains containing the indicated mutations and SwrA overexpressed by IPTG induction (*swrA^+++^*, black circles). The following strains were used to generate this panel: *site1* (DB164, DB198), *site 2* (DB102, DB197), and *site 1+2* double mutant (DB49, DB58). B) Quantitative swarm expansion assays of strains with the indicated sequence between the DegU binding site repeat elements (gray circles), and strains containing the indicated sequence and SwrA overexpressed by IPTG induction (*swrA^+++^*, black circles) The following strains were used to generate this panel: *6A* (DB48, DB196), *6T* (DB47, DB228), *6G* (DB165, DB199), and *6C* (DB166, DB200). C) Quantitative swarm expansion assays of strains in which the 5-8-5 DegU binding site motif was moved farther away (−20 or −10 bp, respectively) or towards (+10 or +20 bp, respectively) the *P_flache_* promoter (gray circles), and strains containing the indicated relocation and SwrA overexpressed by IPTG induction (*swrA^+++^*, black circles) The following strains were used to generate this panel: *-20* (DB488, DB524), *-10* (DB487, DB523), *+10* (DB195, DB220) and *+20* (DB486, DB522). Each data point is the average of three replicates.

We noted that the 5-8-5 inverted repeat was part of the consensus sequence predicted by MEME analysis but the actual output emphasized the thymine-enriched spacer between the repeats rather than the repeats themselves (**Fig S4**). We wondered why the intervening sequence was so highly-conserved and homopolymer replacements were made to test its importance. When six of the spacer residues were changed to either all adenines (A) or all thymines (T), no change on swarming motility was observed and overexpression of SwrA shortened the lag period like wild type (**Fig 6B**). When the residues were changed to all guanines (G), swarming was severely impaired but in a way that could be rescued by SwrA overexpression. When the residues were changed to all cytosines (C) swarming motility was abolished and overexpression of SwrA had little effect. We conclude that the nature of the sequence between the inverted repeats is relevant insofar as only an A-T rich sequence is tolerated for full functionality.

DegU-P bound to the 5-8-5 repeat but did not activate expression until the presence of SwrA widened the region of protection. We wondered whether the effectiveness of the 5-8-5 repeat was dependent on the distance from the −35 box. To test position-dependence, the repeat was moved towards (by deletion) and away (by insertion of randomized sequence) from the promoter by full helical turns of the DNA (10 bp) to maintain register with the promoter (**Fig S7**). Movement of the repeat 20 and 10 bases father away (−20 and −10) abolished and reduced swarming respectively (**Fig 6C**). Overexpression of SwrA however was able to enhance swarming of the −10 reposition. Moving the repeat 10 and 20 bases closer to the promoter (+10 and +20), abolished swarming in a manner that could not be rescued by SwrA overexpression (**Fig 6C**). We conclude that the position of the DegU binding site is important and moving it from its current location impairs swarming motility. Moreover, the strain containing the DegU binding site shifted 20 base pairs closer to the promoter also exhibited mucoid colonies on plates and was difficult to pellet in liquid suggesting additional phenotypic effects. We infer that SwrA expansion of the DegU binding site is likely important to bring the activation complex closer to the promoter, but additional sequence elements may be important for *P_flache_*expression.

The mucoid colony and loose pellet of the strains in which the DegU binding site was moved 20 closer to the promoter was consistent with the phenotype of a SigD mutant, and SigD is encoded at the 3’ end of the *fla/che* operon under *P_flache_*control (11,12,50-52). To measure SigD activity, a reporter in which the SigD-dependent *P_hag_*-promoter was transcriptionally fused to the gene encoding green fluorescent protein (*gfp*) and introduced at an ectopic site in a variety of strains (13). Wild type exhibited a high frequency of brightly fluorescent *P_hag_* ON cells with rare dark *P_hag_* OFF cells **(Fig 7)**. Mutation of either SwrA or DegU decreased the frequency of *P_hag_* ON cells, with OFF cells growing as long chains, whereas mutation of SigD abolished ON cell production and cell chains were ubiquitous (**Fig 7**). Strains in which the DegU binding site was moved 20 or 10 base pairs farther away (+20 and +10) or 10 base pairs closer (−10) to the promoter produced populations similar to that of either the SwrA or DegU mutant, consistent with the inability of the protein complex to activate the *P_flache_* promoter properly **(Fig 7)**. The strain that moved the DegU binding site 20 base pairs closer however, produced a population similar to that of the SigD mutant, indicating a more severe defect in *P_flache_* promoter activity than could be explained simply by impairment of SwrA/DegU **(Fig 7)**. SwrA/DegU did not become inhibitory when the DegU binding site was moved 20 base pairs closer as mutation of either protein failed to restore *P_hag_* expression (**Fig S8**). Alignment of *P_flache_* promoter sequences from closely related species indicated a high degree of conservation including homopolymeric A and T tracts resembling a promoter-enhancing UP element **(Fig 5C, Table S8)** (53–56). We conclude that DegU likely evolved to bind as close to the *P_flache_* promoter as possible while maintaining an UP element constrained to reside immediately adjacent to the −35 box. In order to function as an activator however, an additional factor SwrA was required to expand DegU binding epigenetically closer to the promoter and enhance interaction with RNA polymerase (**Fig 8A**).

**Figure legend 7:**
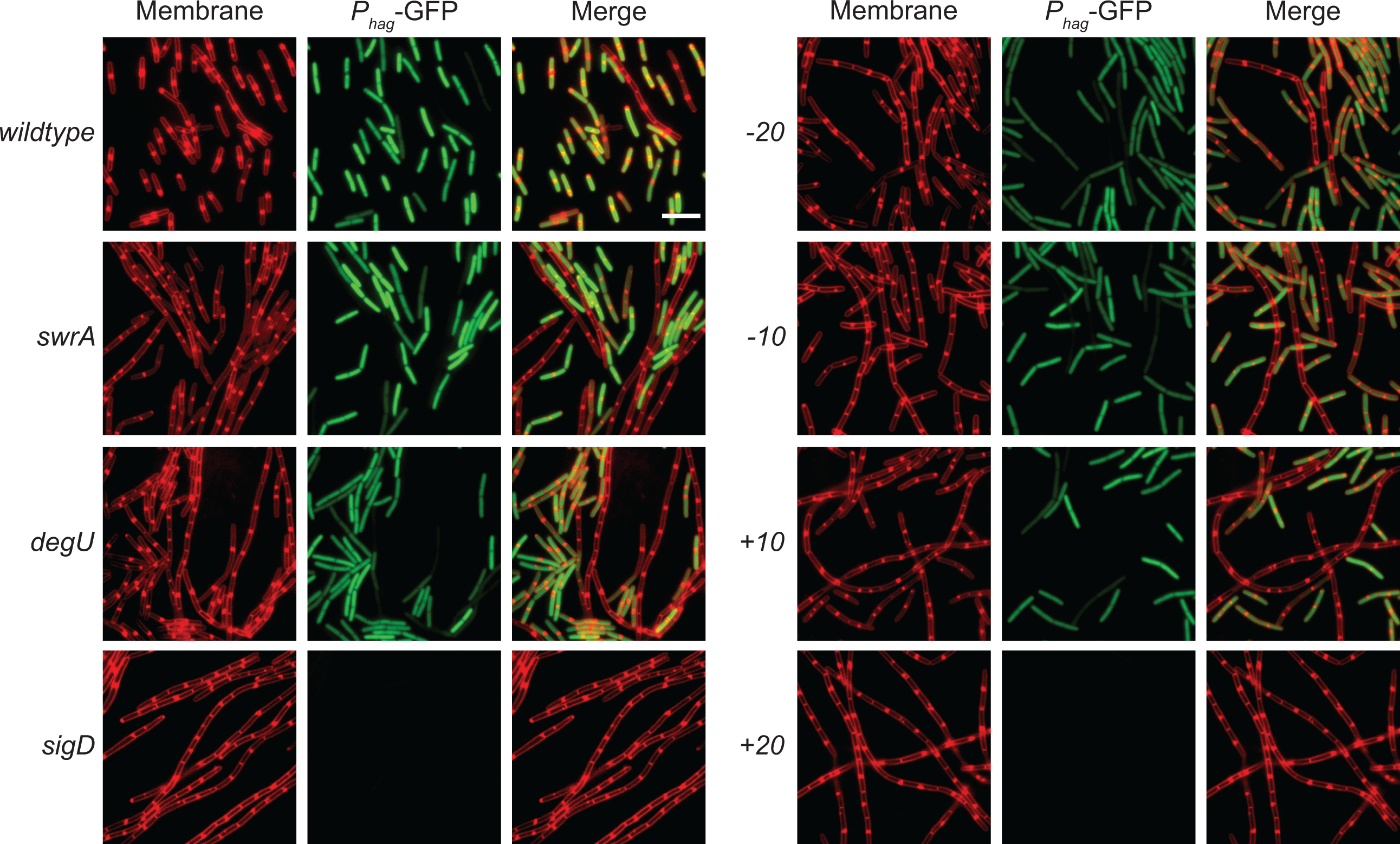
Moving the DegU binding site towards *P_flache_*resembles a *sigD* mutant. Fluorescent micrographs of strains expressing *P_hag_*-GFP in the indicated genetic backgrounds. Membrane was false colored red and *P_hag_*-GFP reporter was false colored green. The following strains are used to generate this panel: *wt* (DK3858), *swrA* (DB549), *degU* (DB550), *sigD* (DB612), −*20* (DB521), *-10* (DB520), *+10* (DB504) and *+20* (DB519). Scale bar is 8µm.

**Figure legend 8.**
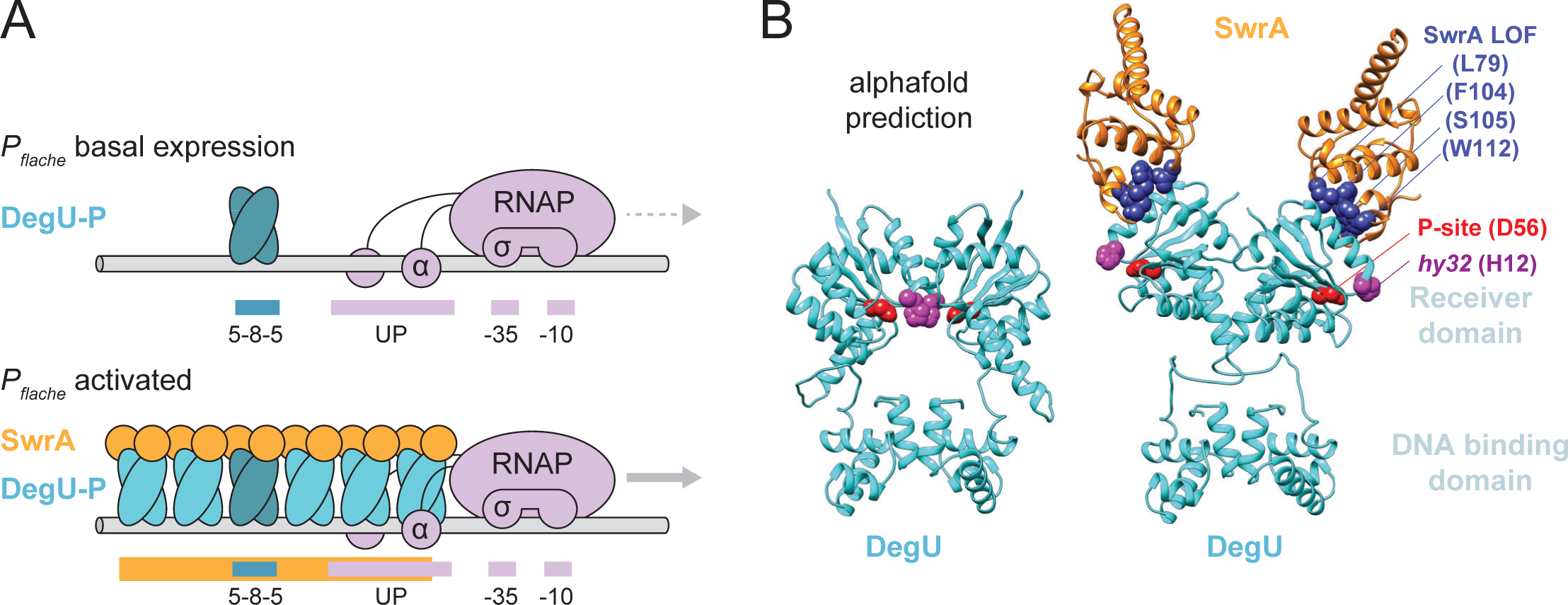
SwrA activates flagellar gene expression by expanding DegU over an UP element. A) Model of SwrA inducing DegU oligomerization. Top, a dimer of DegU-P (blue) binds to the *P_flache_*promoter region (gray cylinder) at the 5-8-5 inverted repeat site. Meanwhile, RNA polymerase (pink) α subunit and σ subunit bind at the UP element and −35/-10 boxes respectively to initiate a basal level of expression (dashed gray arrow). Bottom, SwrA binds to DegU-P causing it to oligomerize and make contact with the α C-terminal domain and enhance expression (solid gray arrow). B) AlphaFold-Multimer prediction of the DegU dimer (cyan) and DegU dimer bound by SwrA (orange). Red, residue phosphorylated by DegS. Purple, residue mutated by the hyperactive *hy32* allele. Dark blue, residues identified in a forward genetic screen which when mutate abolished SwrA activity (16). AlphaFold-Multimer prediction confidence graphs included in **Fig S10**.

## DISCUSSION

SwrA is a poorly-understood transcriptional activator, which while conserved in close relatives of *Bacillus subtilis*, is mutated in most commonly-used laboratory strains (9,40,57-59). How SwrA activates gene expression is unknown but recent work has indicated that SwrA modifies the function of the response regulator DegU (19,20,42). Here we took a global ChIP-seq approach to show that SwrA interacts with DNA indirectly at a subset of DegU binding sites. The DNA binding consensus of DegU is unknown but by focusing on one of the strongest targets, the *P_flache_* promoter, we found an inverted repeat that DegU bound with high affinity. Moreover, addition of SwrA synergized with phosphorylation to enhance DegU DNA binding affinity and caused DegU to protect a region larger than the inverted repeat. We hypothesize that the inverted repeat in *P_flache_* represents the DegU consensus binding site and that SwrA-induced DegU oligomerization that may permit binding at other sites in the chromosome where consensus conservation is poor. Moreover, we suggest that SwrA functions as a co-activator to allow DegU to bind DNA far from the promoter while permitting contact with RNA polymerase.

The paradigm of transcriptional activators was established with the pleiotropic CAP/CRP protein of *E. coli* (60,61). Briefly, CAP binds as a dimer to an inverted repeat sequence (62–64) upstream of a weak promoter and increases RNA polymerase recruitment (65–66) by interaction with the C-terminal domain of α subunit (α-CTD) (67–68). The location of the CAP binding site is important, such that if the CAP site is relocated too far from, too close to, or on the opposite face of DNA from the −35 box, activation is impaired (69–72). Like CAP, DegU is a dimeric, pleiotropic, regulatory protein that binds to an inverted repeat sequence upstream of the *P_flache_* promoter. DegU binding to *P_flache_* however was not sufficient and required an additional protein SwrA to increase expression approximately four-fold. Moreover, moving the DegU binding site towards the *P_flache_* promoter caused a near complete failure of expression, a stronger defect than mere deactivation. Thus, SwrA and DegU not only activates expression of the *P_flache_* promoter but must do so in a way that accommodates a critical cis element that intercedes between the promoter and the binding site.

The *P_flache_* promoter-proximal cis-acting sequence is likely an UP element. UP elements are sequences rich in adenine and thymine bases located immediately upstream of the −35 box of certain promoters (53,54,56), and either increase RNA polymerase stability at the promoter, or control open complex formation to initiate transcription (53,54,73,74). We note that the sequence immediately upstream of *P_flache_*is A-T rich, highly-conserved in related species and very similar to previously-characterized UP elements that enhance the flagella-related *P_hag_*and *P_fliD_* promoters in *B. subtilis* (55,75-77). The failure of the *degU* binding site to stimulate *P_flache_* when moved closer to the −35 box suggests that the SwrA/DegU complex cannot substitute for the UP element and that the two serve different functions in promoting transcription. The presence of DegU/SwrA likely increases affinity of RNA polymerase for the promoter, while the UP element is critical for expression and may function in open complex formation. If this is the case, DegU evolved to bind as close to the promoter as possible while not interrupting the UP element, and SwrA extends the reach of DegU.

We infer that SwrA expands the region of protection on *P_flache_* by changing the oligomeric state of the DegU response regulator. There is no evidence that SwrA interacts with DNA directly and a bacterial two-hybrid assay indicated that SwrA interacted with the N-terminal domain of DegU far from the DNA binding surface (19). The site of DegU-SwrA interaction is supported by AlphaFold-Multimer and SwrA residues essential for activity are located at the interface (**Fig 8B**) (16). Moreover, the AlphaFold model predicts that SwrA binding induces freedom of rotation in the DegU N-terminal dimerization domain, which could provide a mechanism for lateral oligomerization. SwrA-induced oligomerization might also explain the diffuse appearance of some EMSA supershifted bands and why high concentrations of both proteins appeared to cause clogging of polyacrylamide wells. DegU oligomerization might also permit interaction with otherwise weak binding sites and explain why an inverted-repeat consensus binding site was difficult to detect. Finally, we note the enigmatic DegU^hy32^ allele selected in the absence of SwrA sits at the DegU dimerization interface and might induce mobility of the DegU N-terminal domain and partially mimic the SwrA-bound state (**Fig 8B**) (21,22,33,35,42,43).

Why some promoters require UP elements and some do not is unknown. Here, the *P_flache_*promoter requires an UP element for basal expression but is also activated by DegU/SwrA to further increase flagellar gene expression. We posit that a complex transcription factor is needed to accommodate spatial juxtaposition of two incompatible cis-element sequences. While SwrA/DegU may be unique, we note similarities to a number of other poorly-understood activators. For example, RcsA is a small helix-turn-helix protein that modulates the expression of a subset of promoters governed by RcsB to activate polysaccharide biosynthesis (78–80). In addition, Sxy/TfoX is a small protein that modulates a subset of promoters controlled by CAP to activate DNA uptake machinery (81–83). The FlhDC master flagellar regulator of *E. coli/S.enterica* is also bipartite, and while the literature disagrees on which protein(s) binds DNA, productive flagellar gene transcription only occurs in the presence of both (7,84-88). Finally, SwrA, RcsA, Sxy, and FlhC all regulate either long operons or transmembrane nanomachines and each is proteolytically regulated by the Lon protease (17,89-92). Perhaps complex enzymes or machines encoded in long operons require UP elements and consequentially evolved heteromeric transcription factors to activate gene expression in response to environmental input.

## MATERIALS AND METHODS

### Strain and growth conditions

*B. subtilis* and *E. coli* strains were grown in lysogeny broth (LB) (10 g tryptone, 5 g yeast extract, 5 g NaCl per liter) broth or on LB plates fortified with 1.5 % Bacto agar at 37°C. When appropriate, antibiotics were added at the following concentrations: ampicillin 100 μg/ml (amp), kanamycin 5 μg/ml (kan), chloramphenicol 5 μg/ml (cm), spectinomycin 100μg/ml (spec), tetracycline 10 μg/ml (tet), and erythromycin 1μg/ml plus lincomycin 25μg/ml (mls). Isopropyl β-D-thiogalactopyranoside (IPTG, Sigma) was added to LB medium at 1mM concentration when required.

### Strain construction

All PCR products were amplified from *B. subtilis* chromosomal DNA, from the indicated strains. Constructs were transformed into the naturally competent strain DK1042 (3610 *comI^Q12L^*) (93) and transduced using SPP1-mediated generalized transduction to other genetic backgrounds (94).

#### SPP1 phage transduction

To 0.2 ml of dense culture (OD600-0.6-1.0), grown in TY broth (Atoclaved LB broth supplemented with 10 mM MgSO_4_ and 100 µM MnSO_4_), serial dilutions of SPP1 phage stock were added and statically incubated for 15 min at 37 °C. To each mixture, 3 ml TYSA (molten TY supplemented with 0.5% agar) was added, poured atop fresh TY plates, and incubated at 37 °C overnight. Top agar from the plate containing near confluent plaques was harvested by scraping into a 15 mL conical tube, vortexed, and centrifuged at 5,000 X g for 10 min. The supernatant was passed through a 0.45 µm syringe filter and stored at 4 °C. Recipient cells were grown to OD600-0.6-1.0 in 3 ml TY broth at 37 °C. One ml cells were mixed with 25 µl of SPP1 donor phage stock. Nine mL of TY broth was added to the mixture and allowed to stand at 37 °C for 30 min. The transduction mixture was then centrifuged at 5,000 X g for 10 min, the supernatant was discarded, and the pellet was resuspended in the remaining volume. 100 µL of cell suspension was plated on TY fortified with 1.5% agar, the appropriate antibiotic, and 10 mM sodium citrate and incubated at 37 °C overnight. All strains used in this study are listed in **Table 1**. All primers used to build strains for this study are listed in **Table S6** and all plasmids are listed in **Table S7**.

**TABLE 1:**
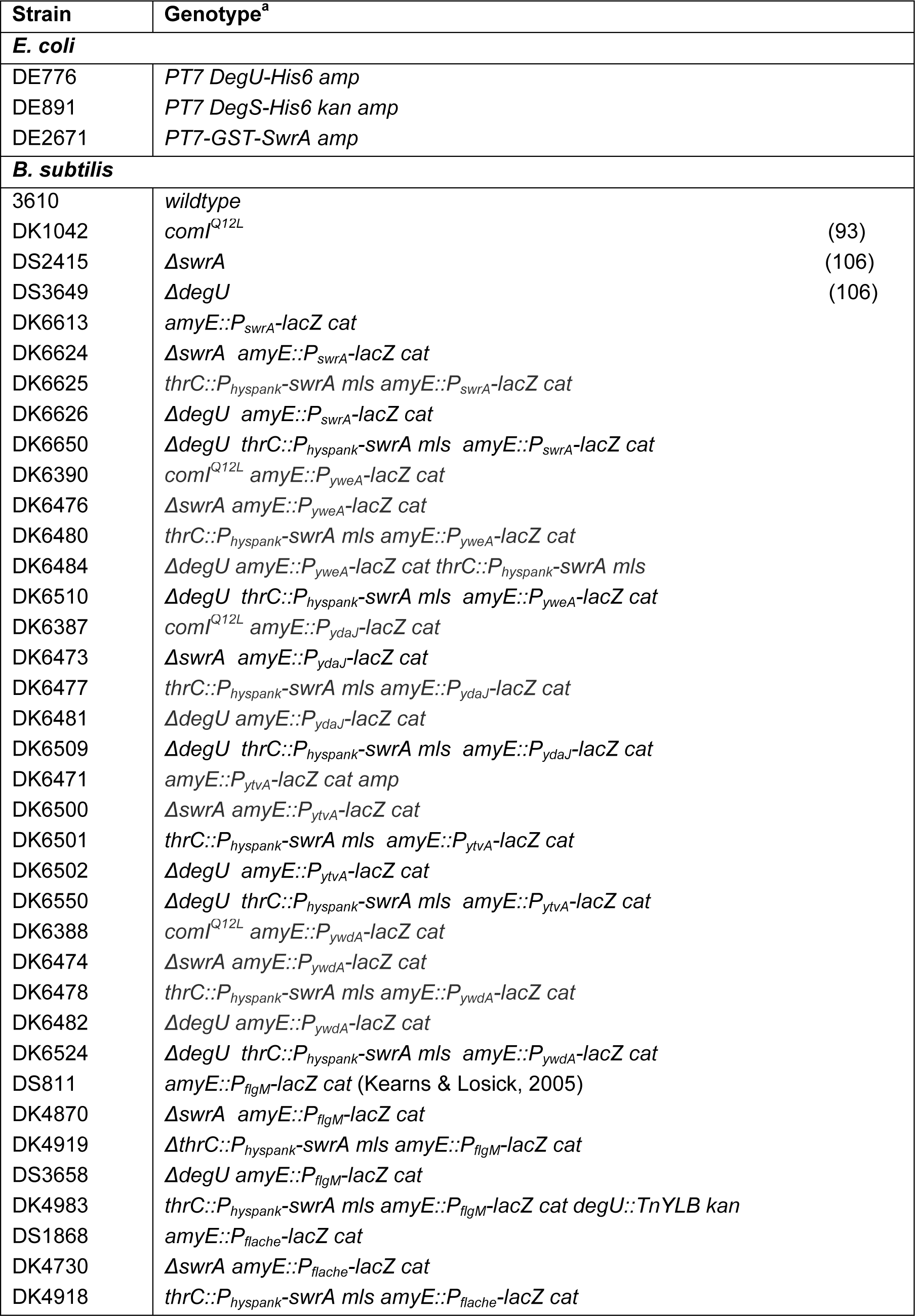
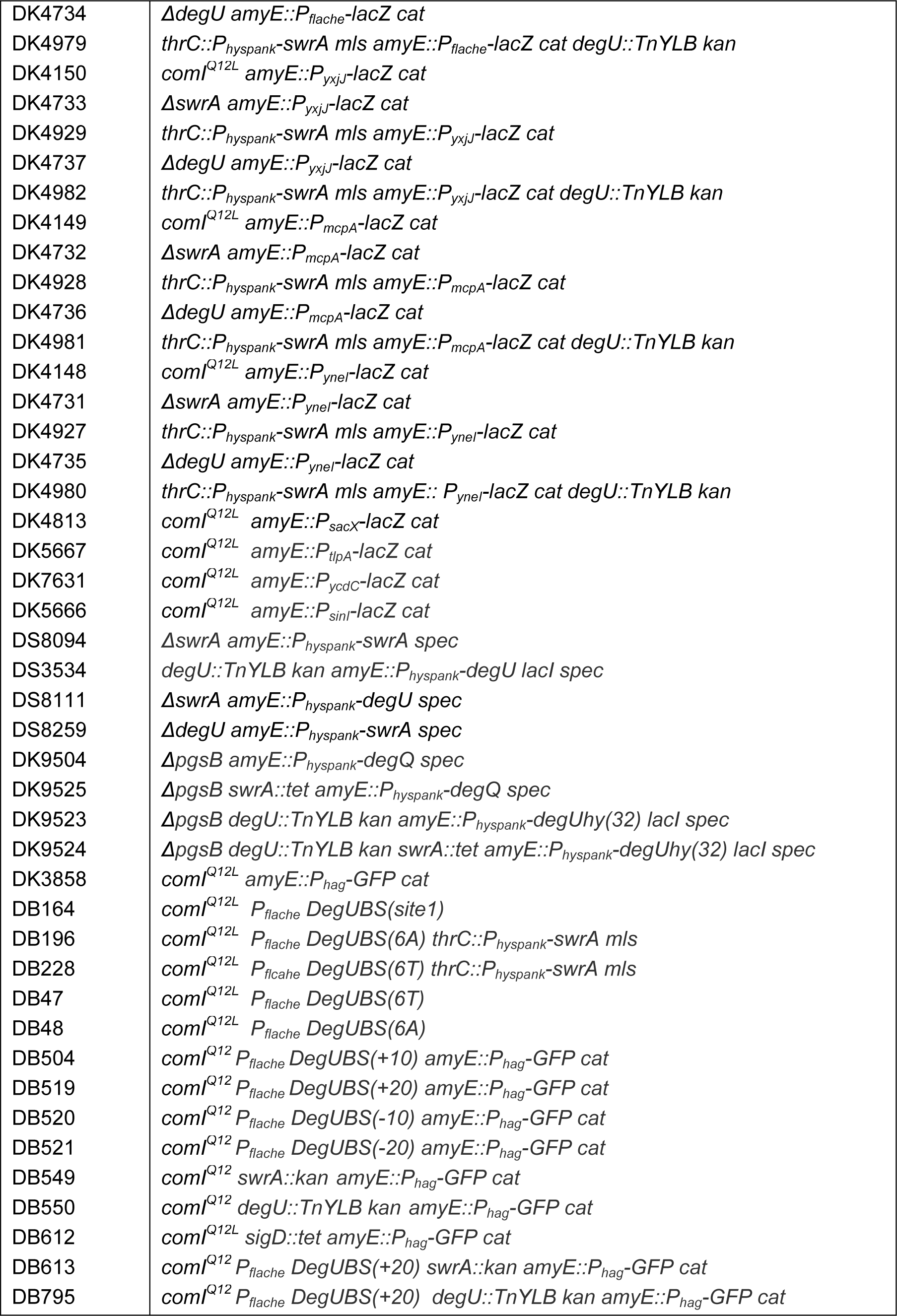
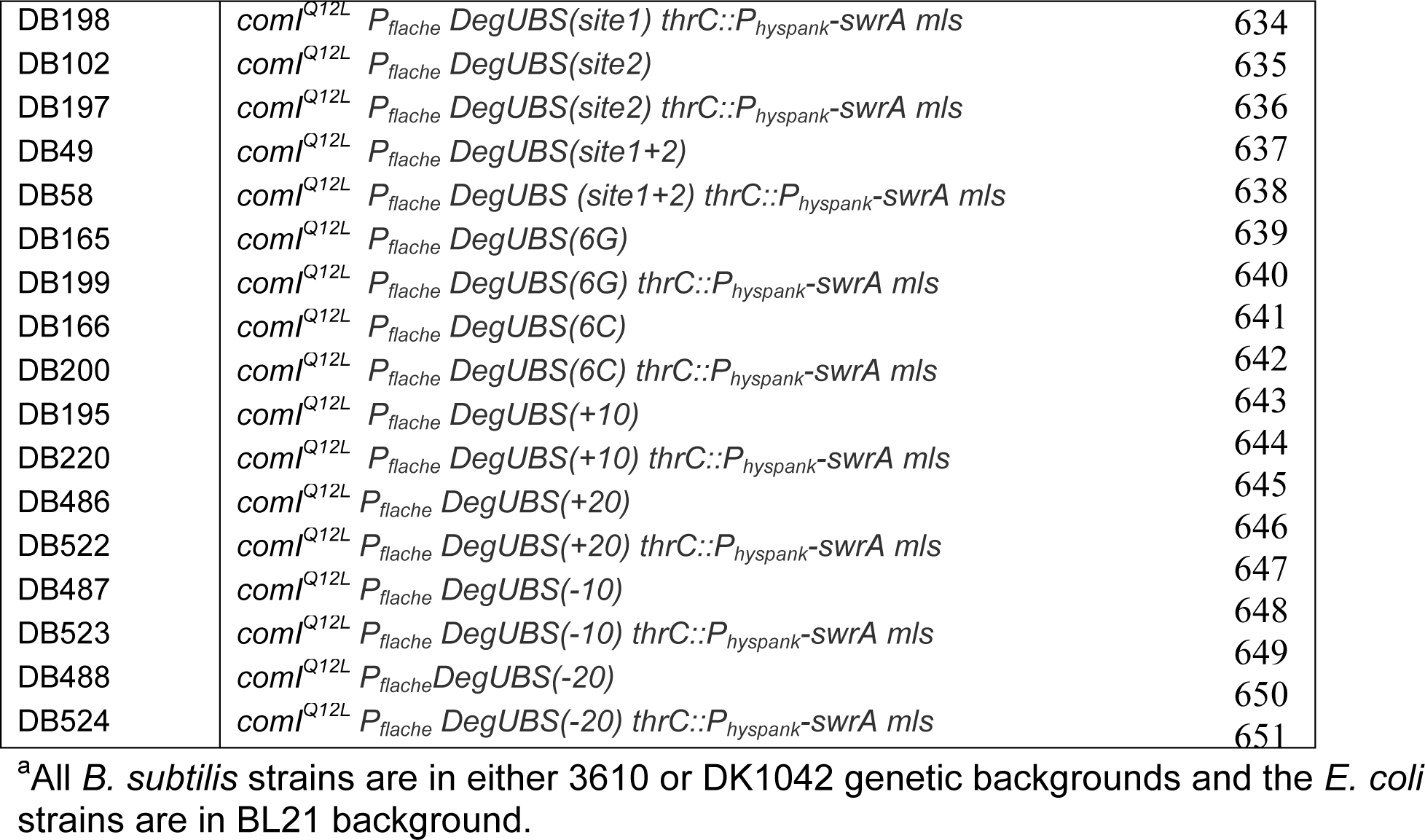
Strains.

#### Transcriptional reporter constructs

The *P_swrA_, P_yxjJ,_ P_mcpA,_ P_yneI,_ P_ydaJ,_ P_ywdA,_ P_sinI_* and *P_tlpA_* promoter regions were amplified from *B. subtilis* strain DK1042 as the template using the primer pairs (353/354), (5120/5121), (5118/5119), (5116/5117), (6481/6482), (6483/6484), (6187/6188) and (6189/6190), respectively. The amplicons were digested with EcoRI and BamHI and ligated into the EcoRI and BamHI sites of pDG268 containing the *lacZ* gene and the *cat* gene for chloramphenicol resistance betweens arms of amyE (95) to generate pDP144, pDP464, pDP463, pDP462, pAM08, pAM09, pDP521 and pDP522.

The *P_ytvA_* and *P_yweA_* promoter regions were amplified from *B. subtilis* strain DK1042 as the template using the primer pair (6487/6488) and (6489/6490) respectively. The amplicons were digested with MfeI and BamHI and ligated into the EcoRI and BamHI sites of pDG268 to generate pAM11 and pAM12.

The *P_sacX_* promoter region was amplified from *B. subtilis* strain DK1042 as the template using the primer pair (6069/6064). The amplicon was digested with HinDIII and BamHI and ligated into the HinDIII and BamHI sites of pDG268 to generate pAM01.

#### Native site mutants

Native site mutants (**Fig S7**) were created by allelic replacement. To generate the following mutations at the native site, the *P_flache_* region was amplified from DK1042 chromosomal DNA using the primer pairs *6T* (7817/7820, 7819/7818), *6A* (7817/7822, 7821/7818), *site1* (7817/7824, 7823/7818), *site2* (7817/7826, 7825/7818), *site1+2* (7817/7828, 7827/7818), *6G* (7817/7870, 7869/7818), *6C* (7817/7872, 7871/7818), *+10* (7817/7887, 7886/7885), *+20* (7817/7980, 7979/7885), *-10* (7817/7982, 7981/7885) and *-20* (7817/7984, 7983/7885). The fragments were assembled by Gibson assembly (96) and the assembled product was digested with EcoRI and SalI. and ligated into the EcoRI and SalI site of pminiMAD carrying a temperature sensitive origin of replication in B. subtilis and the *erm* gene conferring mls resistance (97), to generate pAM39, pAM40, pAM41, pAM42, pAM43, pAM48, pAM49, pAM56, pAM67, pAM68 and pAM69 respectively. The plasmids were passaged individually through *recA+ E. coli* strain TG1, transformed into DK1042 and plated at restrictive temperature for plasmid replication (37°C) on LB agar supplemented with mls to select for transformants with single crossover plasmid integration. Plasmid eviction was ensured by growing the strains for 14 hours at a permissive temperature for plasmid replication (22°C) in the absence of mls selection. Cells were serially diluted and plated on LB agar plates in the absence of mls. Individual colonies were replica patched on LB agar plates with and without mls to identify mls sensitive colonies that have successfully evicted the plasmid. Chromosomal DNA was isolated form the colonies that had excised the plasmid and allelic replacement was confirmed by PCR amplification of the *P_flache_* region using primers 1921 and 3042 followed by sequencing using the same set of primers individually.

### Swarm expansion assay

Cells were grown to mid-log phase (OD_600_ 0.3-1.0) at 37°C in lysogeny broth (LB) and resuspended to and OD_600 of_ 10 in pH 8.0 PBS (137 mM NaCl, 2.7 mM KCl, 10 mM Na_2_HPO_4_, and 2 mM KH_2_PO_4_) containing 0.5% India ink (Higgins). Freshly prepared LB plates fortified with 0.7% bacto agar (25 mL per plate) was dried for 10 min in a laminar flow hood, centrally inoculated with 10 μL of the cell suspension, dried for another 10 min, and incubated at 37 °C. The India ink demarks the origin of the colony and the swarm radius was measured relative to the origin every 30 min. For consistency, an axis was drawn on the back of the plate and swarm radii measurements were taken along this transect. IPTG was added to the medium at final concentration of 1mM to the LB broth and plates when appropriate.

### β-galactosidase assay

*B. subtilis* strains were grown in LB broth at 37°C with constant rotation to OD (0.7-1.0). One ml of cells were harvested and assayed for β-galactosidase activity. The OD_600_ of each sample was measured and the cells were centrifuged and resuspended in 1 ml of Z-buffer (40 mM NaH_2_PO_4_, 60 mM Na_2_HPO_4_, 1 mM MgSO_4_, 10 mM KCl and 38 mM β-mercaptoethanol). To each sample, lysozyme was added to a final concentration of 0.2 mg/ml and incubated at 30°C for 15 minutes. Each sample was diluted appropriately in 500 μl of Z-buffer and the reaction was started with 100 µl of start buffer (4 mg/ml 2-nitrophenyl β-D-galactopyranoside (ONPG) in Z-buffer) and stopped with 250 µl 1 M Na_2_CO_3_. The OD_420_ of the reaction mixtures were recorded and the β-galactosidase specific activity was calculated according to the equation: (OD_420_/time x OD_600_)] x dilution factor x 1000. IPTG was added to the medium at final concentration of 1mM to the LB broth and plates when appropriate. Average β-galactosidase activities and standard deviations in Figure 2 are presented in **Table S5**.

### DegU-His_6_ protein purification

The DegU-His_6_ fusion protein expression vector pNW43 was transformed into BL21(DE3) *E. coli*, and the resulting strain, DE776, was grown to an OD_600_ of 0.7 at 37°C with constant shaking in 1 liter of Terrific Broth (12 g tryptone, 25 g yeast extract, 4ml glycerol and 100 ml sterile potassium phosphate solution (2.31 g KH_2_PO_4_, 12.54 g K_2_HPO_4)_ supplemented with 100 μg/ml ampicillin. Protein expression was induced by adding 1 mM IPTG, and growth was continued overnight at 16°C. Cells were pelleted, resuspended in lysis buffer (50mM Tris HCl pH7.6 and 150mM NaCl), treated with lysozyme and lysed by sonication. Lysed cells were centrifuged at 8000xg for 30 min at 4°C. Supernatant was isolated and combined with 2 ml of nickel-nitrilotriacetic acid (Ni-NTA) His Bind resin (Novagen), equilibrated in lysis buffer and incubated for 1 hour at 4°C with gentle rotation. The resin-lysate mixture was poured into a 1-cm separation column (Bio-Rad), the resin was allowed to pack, and the lysate was allowed to flow through the column. The resin was washed with wash buffer (50mM Tris HCl pH7.6, 150mM NaCl and 5mM Imidazole). DegU-His_6_ protein bound to the resin was eluted in a stepwise manner using lysis buffer supplemented with 10, 30, 100, 250 and 500mM Imidazole. Elution products were separated by 15 % sodium dodecyl sulfate-polyacrylamide gel electrophoresis (SDS-PAGE) and Coomassie stained to verify purification of the DegU-His_6_ protein and fractions containing clean DegU-His_6_ were pooled and concentrated to 2mL. Finally, the concentrated protein was purified via size exclusion chromatography using a Superdex 75 16/60 column (GE Heathcare). DegU-His_6_ was stored in storage buffer (20mM Tris pH7.6, 10% glycerol, 200mM NaCl) at −80C. Concentration of protein was determined by Bradford assay (Bio-rad)

### DegS-His_6_ protein purification

The DegS-His_6_ fusion protein expression vector pYH8 was transformed into BL21(DE3) *E. coli*, and the resulting strain, DE891, was grown to an OD_600_ of 0.5 at 37°C with constant shaking in 500ml of LB Broth supplemented with 25 μg/ml kanamycin. Protein expression was induced by adding 1 mM IPTG, and growth was continued for an additional 4 hours at 30°C. Cells were pelleted, resuspended in lysis buffer (50mM Tris HCl pH7.6 and 150mM NaCl), treated with lysozyme and lysed by sonication. Lysed cells were centrifuged at 8000xg for 30 min at 4°C. Insoluable pellets were resuspended in 4mL of resuspension buffer (6M guanidine HCl, 50mM Tris HCl pH7.6 and 150mM NaCl), stored on ice for 30 mins followed by centrifugation at 30,000xg for 15 minutes. The supernatant was isolated and combined with 2 ml of nickel-nitrilotriacetic acid (Ni-NTA) His Bind resin (Novagen), incubated at room temperature for 20mins and poured into a 1-cm separation column (Bio-Rad). To renature the lysate-resin mixture, the column was serially washed with 10ml of lysis buffer supplemented with 8, 6, 4 and 2M Urea respectively, followed by a final wash of 10ml lysis buffer. DegS-His6 was serially eluted with lysis buffer supplemented with 2ml 100mM, 250mM and 500mM imidazole respectively. Elution products were separated by 15 % sodium dodecyl sulfate-polyacrylamide gel electrophoresis (SDS-PAGE) and Coomassie stained to verify purification of the DegS-His_6_ protein and fractions containing clean DegS-His_6_ were pooled and concentrated to 2mL. Finally, the concentrated protein was dialyzed against storage buffer (50mM Tris-HCl pH7.6, 200mM KCl, 10mM MgCl2, 1mM dithiothreitol (DTT), 20% (v/v) glycerol) and stored at −80°C. Concentration of protein was determined by Bradford assay (Bio-rad).

### GST-SwrA protein purification

The GST-SwrA fusion protein expression vector pSM94 was transformed into BL21(DE3) *E. coli*, and the resulting strain, DE2671, was grown to an OD_600_ of 0.5 at 37°C with constant shaking in 500ml of LB broth supplemented with 100 μg/ml ampicillin. Protein expression was induced by adding 1 mM IPTG, and growth was continued for an additional 4 hours at 30 C. Cells were pelleted, resuspended in lysis buffer (25 mM Tris-HCl pH 8.0, 1 mM DTT, 1 mM EDTA, 0.1% Triton X-100, 150 mM NaCl, and protease inhibitor cocktail (Roche)), treated with lysozyme and lysed by sonication. Lysed cells were centrifuged at 8000xg for 30 min at 4°C. The supernatant was isolated and combined with 2 mL of Glutathione-Sepharose resin (GE Healthcare) and incubated for 1 hour at 4°C. The mixture was poured into a a 1-cm separation column (Bio-Rad) and the column was washed with wash buffer (25 mM Tris-HCl pH 8.0, 1 mM DTT, 0.1% NP-40, 250 mM NaCl, 10% glycerol, and protease inhibitor cocktail (Roche)). GST-SwrA was eluted form the resin using elution buffer 25 mM Tris-HCl pH 8.5, 20 mM glutathione, 1 mM DTT, 250 mM NaCl, 10% glycerol, and protease inhibitor cocktail (Roche)). Elution products were separated by 15 % sodium dodecyl sulfate-polyacrylamide gel electrophoresis (SDS-PAGE) and Coomassie stained to verify purification of the GST-SwrA protein and fractions containing clean GST-SwrA were pooled and concentrated to 2mL. Finally, the concentrated protein was dialyzed against storage buffer (25 mM Tris-HCl [pH 8.0], 1 mM DTT, 250 mM NaCl, 10% glycerol, 20% sucrose) and stored at −80°C. Concentration of protein was determined by Bradford assay (Bio-rad).

### Antibody generation

1 mg His_6_-DegU protein was sent to Cocalico Biologicals for serial injection into a rabbit host for antibody generation. The antibody was purified from serum by mixing it with His_6_-DegU-conjugated Affigel-10 and incubating overnight at 4 °C. The slurry was loaded onto a 1 cm column (BioRad) and eluted with 100 mM glycine (pH 2.5) dropwise and neutralized with 2 M unbuffered Tris base. Elutions were separated by SDS-PAGE and Coomassie stained. Peak elutions were pooled, dialyzed into 1 X PBS, 50% glycerol, and BSA was added to a final concentration of 1 mg/ml prior to storage at −20 °C.

### Chromatin Immunoprecipitation Sequencing (ChIP-Seq)

*Bacillus subtilis* cultures were grown to an OD_600_ of 1.0 at 37°C with constant rotation. 20 ml of cells were cross-linked for 30 minutes at room temperature using 3% formaldehyde (Sigma), quenched with 125 mM glycine, washed with PBS, and then lysed. DNA was sheared to an average fragment size of ∼200 bp using Qsonica sonicator (Q8000R), and then incubated overnight at 4°C with α-SwrA or α-DegU as indicated. Immunoprecipitation was performed using Protein A Magnetic Sepharose beads (Cytiva #45002511), washed, and DNA was eluted in TES (50mM Tris pH8, 10mM EDTA and 1% SDS). Crosslinks were reversed overnight at 65°C. DNA samples were treated with a final concentration of 0.2mg/ml RNaseA (Promega #A7973) and 0.2mg/ml Proteinase K (NEB #P8107S) respectively, and subsequently extracted using phenol/chloroform/isoamyl (25:24:1). DNA samples were then used for library preparation using NEBNext UltraII DNA library prep kit (NEB #E7645L). Paired end sequencing of the libraries was performed using Illumina NextSeq 550 platform and atleast 3 million paired end reads were obtained for each sample. Two or three biological replicates were sequenced for each sample.

### Whole genome sequencing (WGS)

*Bacillus subtilis* cultures were grown to an OD_600_ of 1.0 at 37°C with constant rotation and 5ml of cells were collected, pelleted and DNA was extracted using Qiagen DNeasy kit (#69504). Genomic DNA was sonicated using Qsonica sonicator (Q8000R) and the sonicated DNA was used to prepare libraries using the NEBNext UltraII DNA library prep kit (NEB #E7645L). Paired end sequencing of the libraries was performed using Illumina NextSeq 550 platform and at least 3 million paired end reads were obtained for each sample. Data from WGS was used as input for the ChIP.

### Analysis of ChIP-Seq and WGS data

Sequencing reads for both ChIP and WGS were mapped individually to *B. subtilis* 3610 genome (CP020102) (98) using CLC Genomics Workbench software (Qiagen). The enrichment at ribosomal RNA locations were eliminated and the number of reads mapped to each base pair in the genome was exported into a .csv file. Data was normalized to the total number of reads and fold enrichment was calculated as the ratio of number of reads at each genome location in ChIP-Seq and WGS (ChIP/input). Analysis was performed and graphs were plotted in 1kb bins to show enrichment across the entire genome using custom R-scripts. When required, individual peaks were plotted in 10bp bins across a 4kb range centered around the peak summit. Detailed protocols and scripts are available upon request.

### MEME analysis

200bp sequence surrounding the DegU peak center in *WT, degQ^+++^* and *degU degUhy32^+++^* (**Table S1, S2 and S3**) was extracted using a custom perl script and a fasta file was created. Sequences were subjected to Multiple Em for Motif Elicitation (MEME) v 5.5.2 using parameters (meme sequences.fa -dna -oc . -nostatus -time 14400 -mod anr -nmotifs 3 - minw 6 -maxw 30 -objfun classic -revcomp -markov_order 0) (54) (**Table S2**). 30bp highly enriched motif sequences were extracted and sequence logo presented **Fig S4** was created by WebLogo using default parameters (99).

### Electromobility shift assay

DNA probes of 150-200bp regions surrounding DegU ChIP-Seq peak centers were generated by PCR amplification using DK1042 chromosomal DNA and the following primer pairs: *P_flache_*(7231/1782), *P_yxjJ_* (7463/7464), *P_flgM_* (7459/7460), *P_mcpA_* (7554/7555), *P_swrA_* (7461/7462), *P_yneI_* (7465/7466) and *P_comK_* (7229/7230). When required the *P_flache_ (site1+2)* region was amplified using primer pirs (7231, 1782) using DB49 chromosomal DNA as the template. 25nM of DNA was radiolabeled using T4-Polynucleotide kinase (NEB M0201S) and 0.5μl of ATP(γ-^32^P) (Perkin, 3000Ci/mmol) in 20μl reactions. Excess unincorporated ATP(γ-^32^P) was removed by passing the reaction through as G-50 Micro Columns (Cytiva) and the radiolabeled DNA was stored at 4°C until further use. When appropriate, DegU phosphorylation reactions were performed by adding DegU-His_6_ and DegS-His_6_ at a ratio of 1:5 and 1mM cold ATP to kinase buffer (50 mM Tris-HCl (pH 7.6), 0.1 mM EDTA, 10 mM MgCl_2_, 1 mM DTT, 50 mM KCl, 0.1 mg/ml BSA, 10% glycerol) and the reaction was incubated at room temperature for 20 minutes. 20 μl binding reactions were prepared with various concentrations of DegU, DegU-P and SwrA as indicated, 1ul of radiolabeled probe and 5ng/ul polydI-dC (Roche) in binding buffer (50 mM Tris-HCl (pH 7.6), 0.1 mM EDTA, 10 mM MgCl_2_, 1 mM DTT, 50 mM KCl, 0.1 mg/ml BSA, 10% glycerol, 0.1mM ATP) and incubated at 30°C for 30 minutes. 6.5% native polyacrylamide gel was prepared using 19:1 acrylamide/bisacrylamide (Biorad), 1X Tris-Glycine-EDTA buffer (25mM Tris base, 250mM Glycine and 1mM EDTA pH 8.0) and 5% glycerol. Glycerol was added to the binding reaction at a final concentration of 10% to facilitate loading and 12ul of the reaction mixture was fractionated on a 6.5% native gel in 1X TGE running buffer at room temperature for 1 hour at 100V (constant). Gels were dried, exposed to a phosphorimager screen overnight and radioactive signal was detected using Typhoon FLA 9500. Fiji v 2.1.0 (100) was used to quantify the fraction of unbound DNA the fraction of bound DNA was calculated by subtracting the fraction of unbound DNA from 1. DNA binding curves were generated in GraphPad Prism 9 using Non-linear regression and one site specific binding parameters.

### DnaseI-footprinting assay

300−350bp fluorescently labelled DNA probes surrounding the DegU ChIP peak was generated by PCR amplification using a forward primer with a 5′-FAM fluorescent tag (IDT) and a reverse primer with a 5′-HEX fluorescent tag (IDT). *P_flache_* probe was generated using primer pair 7548/7549 and *P_yxj_* probe was generated using primer pair 7562/7563 and DK1042 chromosomal DNA as template. The optimal concentration of DnaseI (NEB #M0570) at which uniform cleavage was observed across the probe was assessed over a range of concentrations (1X – 1024X) prepared in the presence of 1X DNAseI buffer. DegU was phosphorylated as described earlier. 20ul binding reactions were set up in 1X binding buffer to which 5ng/μl polydIdC, 20nM DNA probe, 1X DnaseI buffer and either no protein or indicated amounts of DegU-P and SwrA were added and incubated at 30°C for 30 minutes. 5μl of optimized DNAseI dilution was added to the reaction and incubated at room temperature for 15 minutes. Reaction was quenched by addition of 25μl of 0.5M EDTA pH 8.0. DNA fragments were cleaned using Qaigen MinElute PCR purification kit (#28004). Fragment analysis was performed by Genewiz, Azenta Life Sciences using a 3730 DNA analyzer and fragment size was determined using a GeneScan 500LIZ DNA size standard. Data was analyzed using Peak Scanner software v1.0 and the peak height corresponding each fragment size was exported into a text file. The values were plotted using a custom R-script and DNaseI protection was determined by an absence of peaks across a range of consecutive fragment sizes.

### Alignment of *P_flache_* region

All bacteria that contain both SwrA and DegU were identified by BLAST+ v 2.12.0 (101). *P_flache_* sequence ranging from +1 to −120 (**Fig 5**) and −121 to −250 (**Fig S6**) were extracted and aligned by Clustal Omega v 1.2.4 using default parameters (102). Alignment was shaded using Jalview v 2.11.2.7(103) using a 60% identity threshold and false colored using Adobe illustrator.

### Microscopy

For microscopy, 3ml of LB broth was inoculated with a single colony and grown at 37C to OD600 05-0.8. 1ml of culture was pelleted and resuspended in 30μl 1X PBS buffer containing 5 µg/ml FM 4-64 (Invitrogen #T13320) and incubated for 2 min at room temperature in the dark. Excess dye was washed with 1mL of PBS, cells were spun down and resuspended in a final volume of 30μl of PBS. Flat agarose pads (1% agarose in PBS) were created by on a slide, 5μl of sample was spotted on the Agarose pads and covered with a glass coverslip. Fluorescence microscopy was performed with a Nikon 80i microscope with a phase contrast objective Nikon Plan Apo 100X and an Excite 120 metal halide lamp. FM4-64 was visualized with a C-FL HYQ Texas Red Filter Cube (excitation filter 532-587 nm, barrier filter >590 nm). GFP was visualized using a C-FL HYQ FITC Filter Cube (FITC, excitation filter 460-500 nm, barrier filter 515-550 nm). Images were captured with a Photometrics Coolsnap HQ2 camera in black and white using NIS elements software and subsequently false colored and superimposed using Fiji v 2.1.0.

### Structure prediction

Structure prediction was performed using Alphafold (104). The *B. subtlis* 3610 DegU and SwrA sequences were separated a colon (:) whenever necessary and prediction was performed using parameters colabfold_batch –num-recycle 20 –amber – templates –model-type alphafold2_multimer_v2. The structures were visualized using UCSF Chimera v 1.15 (105).

### Data availability

ChIP-seq and WGS data have been submitted to the Gene Expression Omnibus and we are awaiting accession numbers (accession no. GSExxxx).

## ACKNOWLEDGEMENTS

We thank members of the Kearns lab for helpful discussions, members of the Van Kessel lab for biochemical support and we thank Zhongqing Ren for assistance with ChIP-seq. We thank the Indiana University Center for Genomics and Bioinformatics for high throughput sequencing and AlphaFold-Multimer was performed using IU Carbonate supported in part by Lilly Endowment, Inc., through its support for the Indiana University Pervasive Technology Institute. Support for this work comes from National Institutes of Health R35GM145299 to DZR, R01GM141242, R01GM143182, R01AI172822 to XW, and R35GM131783 to DBK. This research is a contribution of the GEMS Biology Integration Institute, funded by the National Science Foundation DBI Biology Integration Institutes Program, Award #2022049 to XW.

